# Structure–activity relationship of antimycin A-like compounds as photosystem II inhibitors

**DOI:** 10.64898/2026.06.22.733872

**Authors:** Ko Imaizumi, Masatoshi Murai, Hideto Miyoshi, Kentaro Ifuku

## Abstract

Antimycin A (AA) is widely used as an inhibitor of the mitochondrial respiratory chain, targeting the Q_i_ site of cytochrome *bc*_1_ (complex III). In photosynthetic organisms, AA is also well known to inhibit the photosynthetic PROTON GRADIENT REGULATION 5 (PGR5)-dependent cyclic electron flow around photosystem I (CEF-PSI). Although AA is frequently used as a specific inhibitor of PGR5-dependent CEF-PSI in photosynthetic reactions, we recently clarified that some of the major components of AA, which is typically a mixture of closely related compounds, also exert direct inhibitory effects on photosystem II (PSII). Nevertheless, the binding site and binding mode of AA in PSII remain largely unexplored. Structurally, AA consists of a salicylic acid moiety connected via an amide bond to a hydrophobic dilactone ring moiety. To identify important structural factors of AA for exhibiting inhibitory effects on PSII (assessed by Q_A_^−^ reoxidation measurements), we here investigated the relationship between structure and inhibitory potency using 38 AA-like compounds (AALCs), including commercial compounds and a series of synthetic AA analogs. Some AALCs exhibited substantially stronger impacts on PSII than natural AA. High acidity of the phenolic OH and the presence of a free amide NH of the salicylamide moiety were critical for the effects on PSII. In contrast, while the dilactone ring moiety also affected the inhibitory activity, this was replaceable with certain hydrophobic structures. Based on our results, together with the known structure–activity relationship and binding mode of AA in complex III, we propose tentative binding models for AA in PSII.

**Highlights:**

- Structure–activity relationship of AA-like compounds on PSII is examined
- Several AA-like compounds more potent than AA against PSII are identified
- Phenolic OH acidity and free amide NH of salicylamide moiety are key for AA effects
- The dilactone ring moiety is replaceable with certain hydrophobic structures
- Tentative binding models for AA in PSII are proposed

## 1. Introduction

Photosynthetic electron transport proceeds through linear electron flow (LEF) and cyclic electron flow around photosystem I (CEF-PSI). LEF initiates at photosystem II (PSII), a pigment–protein complex that catalyzes light-driven water oxidation at its donor side and reduces plastoquinone (PQ) molecules at its acceptor side (Shen, 2015; Shevela et al., 2023; Imaizumi and Ifuku, 2025a). Light energy absorbed by PSII excites P680, the reaction center chlorophylls of PSII, and charge separation occurs. The electron is transferred from pheophytin (Pheo_D1_) to the non-exchangeable quinone Q_A_ and further to the exchangeable electron acceptor quinone Q_B_, whereas the electron hole drives water oxidation at the Mn_4_CaO_5_ cluster. Double-reduced Q_B_ dissociates from PSII through the plastoquinone exchange channels and enters the PQ pool in the thylakoid membrane. The electrons are then passed to cytochrome (Cyt) *b*_6_*f*, plastocyanin (or Cyt *c*_6_), PSI, and ferredoxin (Fd), and are ultimately used by Fd-NADP^+^ reductase (FNR) to reduce NADP^+^ to NADPH. In contrast, CEF-PSI does not involve PSII, and electrons are cycled from reduced Fd back into the PQ pool. Two well-known CEF-PSI pathways have been proposed. One of them requires the protein PROTON GRADIENT REGULATION 5 (PGR5), while the other is mediated by the chloroplast NADH dehydrogenase-like (NDH) complex (Shikanai, 2024). There has been substantial progress in understanding the structural basis and molecular mechanisms of NDH-dependent CEF-PSI (Ifuku et al., 2011; Peltier et al., 2016; Laughlin et al., 2020; Shen et al., 2022; Su et al., 2022; Introini et al., 2025; Shikanai et al., 2025); however, those of PGR5-dependent CEF-PSI remain largely elusive.

Antimycin A (AA) is an antibiotic produced by *Streptomyces* species (Dunshee et al., 1949). The typically used “AA” is a mixture of closely related compounds, majorly AA1, AA2, AA3, and AA4 (Lockwood et al., 1954; Kluepfel et al., 1970; Schilling et al., 1970) (**Fig. S1**). AA is generally known as an electron transport inhibitor targeting Cyt *bc*_1_ (complex III) in the mitochondrial respiratory chain (Ahmad et al., 1950; Slater, 1973). It specifically binds to the quinone Q_i_ site (quinone reduction site) of complex III, and its binding properties have been extensively investigated through examining the relationship between chemical structure and inhibitory activity (Dickie et al., 1963; Rieske et al., 1967; Miyoshi et al., 1991; Tokutake et al., 1993; Tokutake et al., 1994; Miyoshi et al., 1995; Machida et al., 1999) and structural biology works (Zhang et al., 1998; Gao et al., 2003; Huang et al., 2005; Esser et al., 2008). AA consists of an essential salicylic acid moiety linked via an amide bond to a hydrophobic dilactone ring moiety (van Tamelen et al., 1961; Rieske, 1976). In the field of photosynthesis, AA has also long been recognized as an inhibitor of a CEF-PSI pathway (Tagawa et al., 1963), referred to as the “AA-sensitive pathway” (Labs et al., 2016). Later studies have determined that the PGR5-dependent CEF-PSI pathway, but not the NDH-dependent pathway, is the “AA-sensitive pathway” (Joët et al., 2001; Munekage et al., 2002; Munekage et al., 2004; Sugimoto et al., 2013). Although the mechanism of its inhibition by AA remains unknown (Wang et al., 2018; Rühle et al., 2021; Sarewicz et al., 2021), AA has been widely used as an inhibitor of PGR5-dependent CEF-PSI as if it were a specific inhibitor.

Besides the inhibition of complex III and PGR5-dependent CEF-PSI, AA had also been suggested to affect PSII (Hind, 1968; Cramer et al., 1971; Cramer and Böhme, 1972; Katoh, 1972; Satoh and Katoh, 1972; Yerkes and Crofts, 1992; Miyake et al., 1995); however, this has often been overlooked. Recently, we have reinvestigated the effects of AA on PSII in green plants, and revealed that AA can have direct strong inhibitory impacts on PSII (Takagi et al., 2019; Imaizumi et al., 2025; Imaizumi and Ifuku, 2025b). AA suppressed the oxygen-evolving activity of PSII, perturbed the electron transfer within PSII, and made PSII hypersensitive to light under the presence of inhibitors binding to the Q_B_ site, such as 3-(3,4-dichlorophenyl)-1,1-dimethylurea (DCMU). Furthermore, we observed clear differences in the effects on PSII among the four major AA components; AA1 and AA2 exhibited inhibitory effects on PSII, whereas AA3 and AA4 did not (Imaizumi et al., 2025; Imaizumi and Ifuku, 2025b). This suggested that the binding of AA to PSII is specific, rather than non-specific, and that its effects on PSII are remarkably affected by slight differences in the chemical structures of AAs. Here, we used 38 different AA-like compounds (AALCs) and investigated their effects on PSII (**Fig. 1**). The results give insight into the chemical structures/properties required for the AA-effects on PSII.

**Fig. 1.**
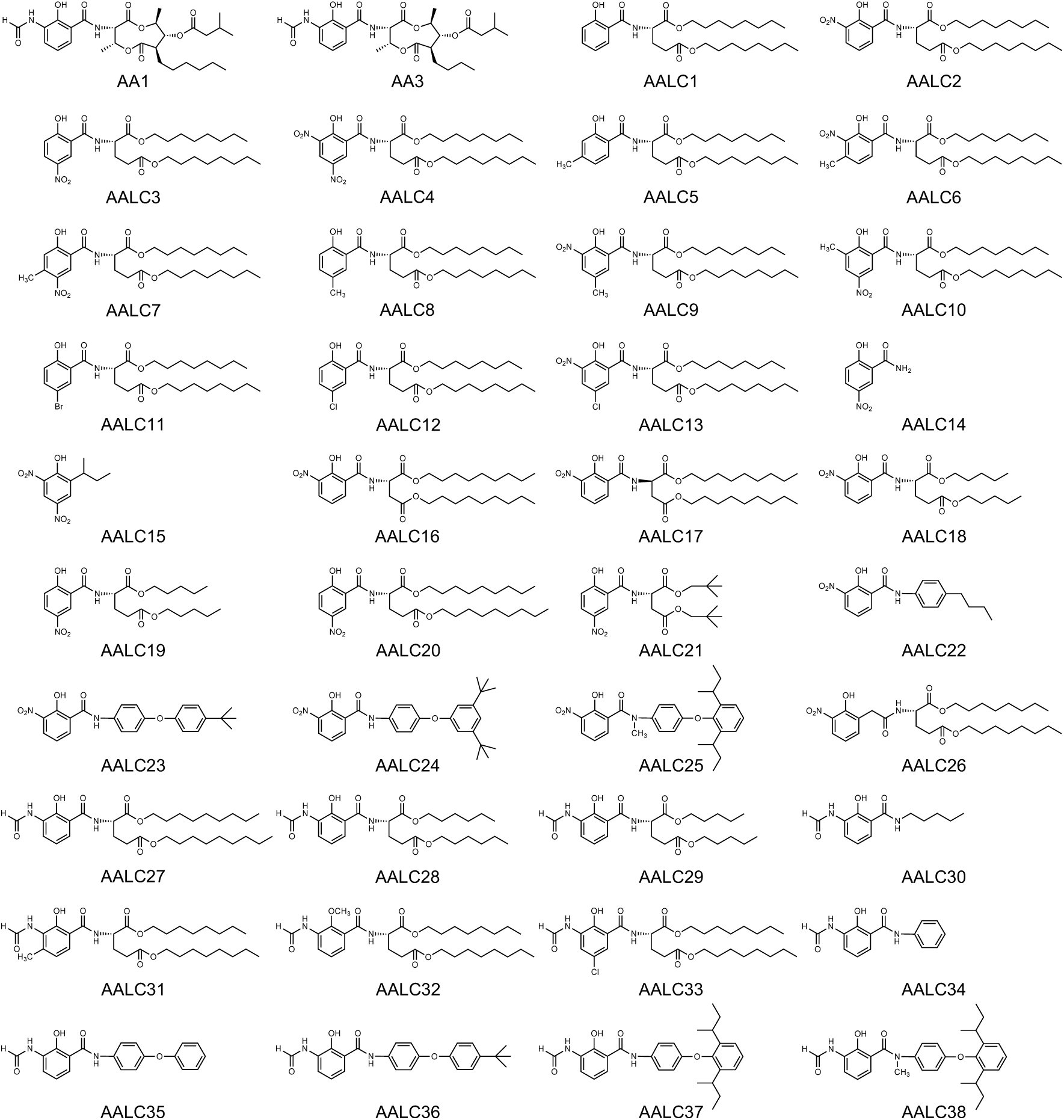
Chemical structures of AA1, AA3, and AALCs. AA1 and 38 AALCs were used in this study. AA3 is shown for reference.

## 2. Materials and methods

### 2.1. Measurement of Q_A_^−^ reoxidation kinetics

Thylakoid membranes were isolated from market spinach with a previously reported method (Imaizumi et al., 2025), modified from Imaizumi et al. (2022). They were suspended in buffer A (20 mM HEPES-NaOH, 1 M betaine, 10 mM NaCl, 5 mM MgCl_2_, pH 7.6) to a final chlorophyll (Chl) concentration of 5 µg Chl mL^−1^, and incubated with inhibitor compounds (AA1, AALCs, and/or DCMU; final concentration of 10 µM) for 3 min at room temperature in complete darkness. Then, the rise of chlorophyll fluorescence induced by a single-turnover flash and its subsequent decay was monitored using the double-modulation fluorometer FL3500 (Photon Systems Instruments) as described previously (Reifarth et al., 1997; Vass et al., 1999; Imaizumi et al., 2025). Data were normalized and fit to a tri-exponential decay equation (Reifarth et al., 1997) using the software OriginPro. For data in the presence of DCMU, in cases where the tri-exponential decay model failed or yielded non-physical parameters, a bi-exponential model was used instead. For data in the absence of DCMU, all three independent replicates for each condition were combined and fitted simultaneously to obtain stable parameter estimates for all components.

### 2.2. *F*_v_/*F*_m_ measurement

Thylakoid membranes (5 µg Chl mL^−1^) suspended in buffer A were incubated with inhibitor compounds for 2 min at room temperature in complete darkness. *F*_v_/*F*_m_ was measured using AquaPen-C AP-C 100 (Photon Systems Instruments).

### 2.3. AA and AA-like compounds

Antimycin A1 was purchased from BioAustralis Fine Chemicals (Batch: AC33.74). 2-Hydroxy-5-nitrobenzamide was purchased from BLDpharm (Lot: ALY787). Dinoseb (2-*sec*-butyl-4,6-dinitophenol) was purchased from FUJIFILM Wako Pure Chemical Corporation (Lot: ESF1914). All other AALCs are the same samples as those used in previous reports (Tokutake et al., 1993; Tokutake et al., 1994; Miyoshi et al., 1995). All inhibitors were dissolved in ethanol and added to the reaction buffer (buffer A) to give a final concentration of 10 µM.

## 3. Results

### 3.1. Effects of co-treatment of AA-like compounds and DCMU on the Q_A_^−^ reoxidation kinetics

We have recently shown that AA has direct inhibitory effects on PSII activity and electron transport within PSII (Takagi et al., 2019; Imaizumi et al., 2025; Imaizumi and Ifuku, 2025b). Measurement of the Q_A_^−^ reoxidation kinetics of thylakoid membranes was found to be effective for investigating the direct effects of AA on PSII in detail (Imaizumi et al., 2025). In this assay, the reduction of Q_A_^−^ by a single-turnover flash leads to a rise in fluorescence yield, and by monitoring the subsequent fluorescence decay in the dark, which reflects Q_A_^−^ reoxidation, we can obtain information on the electron transfer within PSII (Reifarth et al., 1997). In the presence of DCMU, Q_A_^−^ reoxidation proceeds via charge recombination with the donor side of PSII. In contrast, co-treatment with DCMU and AA leads to a significant rise in the final, time-independent, residual fluorescence level (Imaizumi et al., 2025). The amplitude of this residual component was found to be a useful indicator of the AA-effects on PSII. The residual amplitude increased under the presence of AA1 or AA2, while it was not affected by AA3, AA4, or myxothiazol (a complex III Q_o_ site inhibitor). Under the presence of AA mixtures, the rate of increase was dependent on the AA1-plus-AA2 to AA3-plus-AA4 ratio in the mixture.

We conducted Q_A_^−^ reoxidation measurements of thylakoid membranes co-treated with DCMU (10 µM) and either AA1 (10 µM) or AALCs (10 µM), and compared the amplitude of the residual component (**Fig. 2** and **Fig. 3**). AA1 increased the residual amplitude from 7.8% (with only DCMU) to 26.3% (with DCMU and AA1), similar to our previous results (Imaizumi et al., 2025). The effects of AALCs in the presence of DCMU varied greatly. AALC2, 3, 9, 10, 13, 23, and 24 showed very strong effects with residual amplitudes of 45% to 60%. AALC16, 17, 19, and 20 also showed effects stronger than AA1 with residual components between 32% and 38%, whereas the effects of AALC6, 7, 36, and 37 were similar to that of AA1 (residual components between 25% and 28%). AALC15, 18, 21, 22, and 33 induced an increase in the residual component, but to a lesser extent than AA1 (residual amplitudes of 14% to 19%). Addition of AALC4 strongly affected PSII, but a reliable decay curve could not be obtained due to a low signal-to-noise ratio (**Fig. S2a**). The other AALCs did not induce a significant rise in the residual amplitude.

**Fig. 2.**
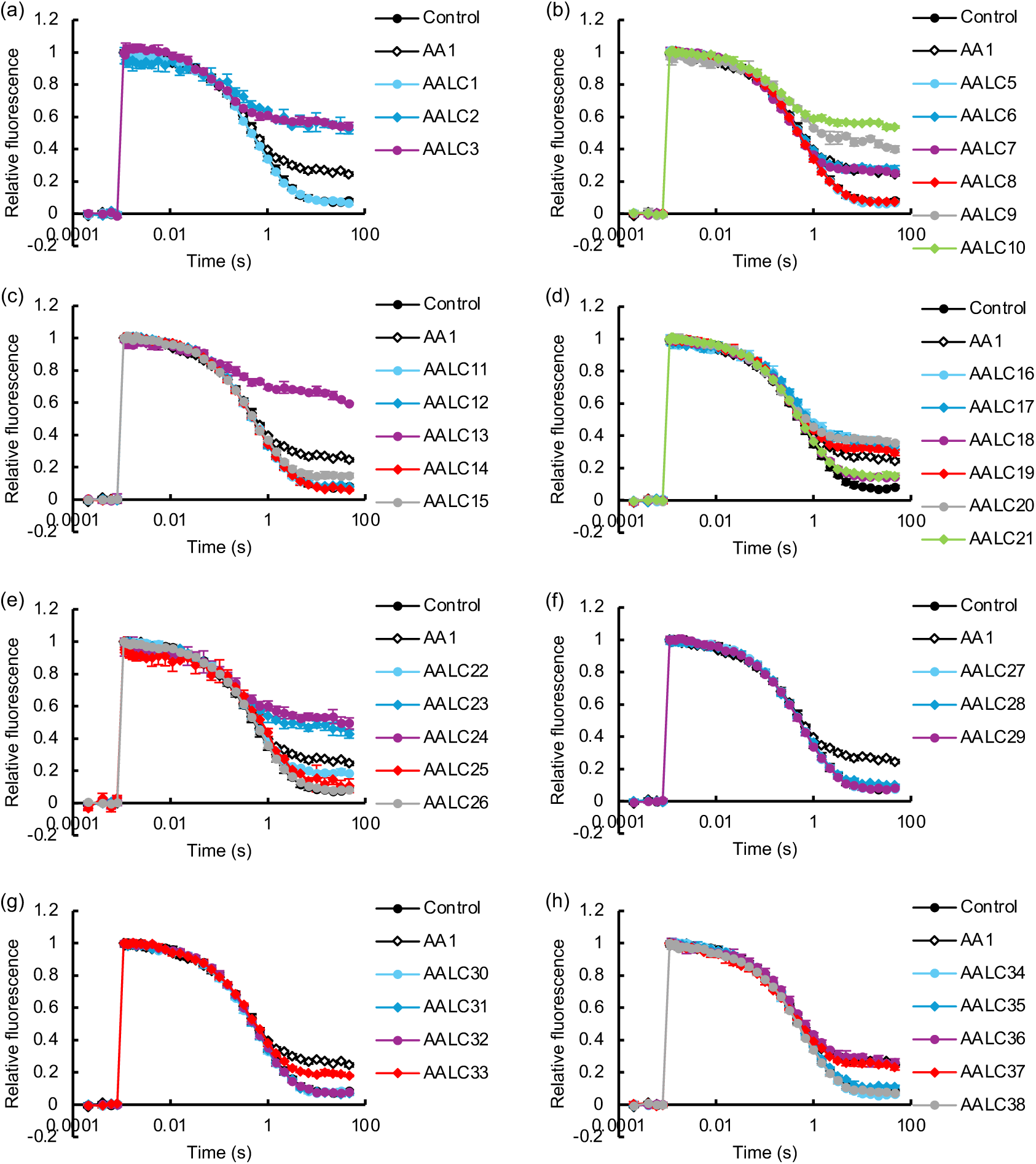
Effects of AALCs on the Q_A_^−^ reoxidation kinetics of PSII in the presence of DCMU. Ethanol (“control”; black filled circles), 10 µM AA1 (black open diamonds), and 10 µM AALCs (colored filled symbols) were added together with 10 µM DCMU to spinach thylakoid membranes. Although all compounds (and the control) were measured in each independent experiment, for clarity, the results for AALCs are presented in eight separate panels (**a**–**h**), with the same control and AA1-treated data shown in all panels. Data are mean ± SD (*n* = 3). In some instances, error bars are smaller than the symbols.

**Fig. 3.**
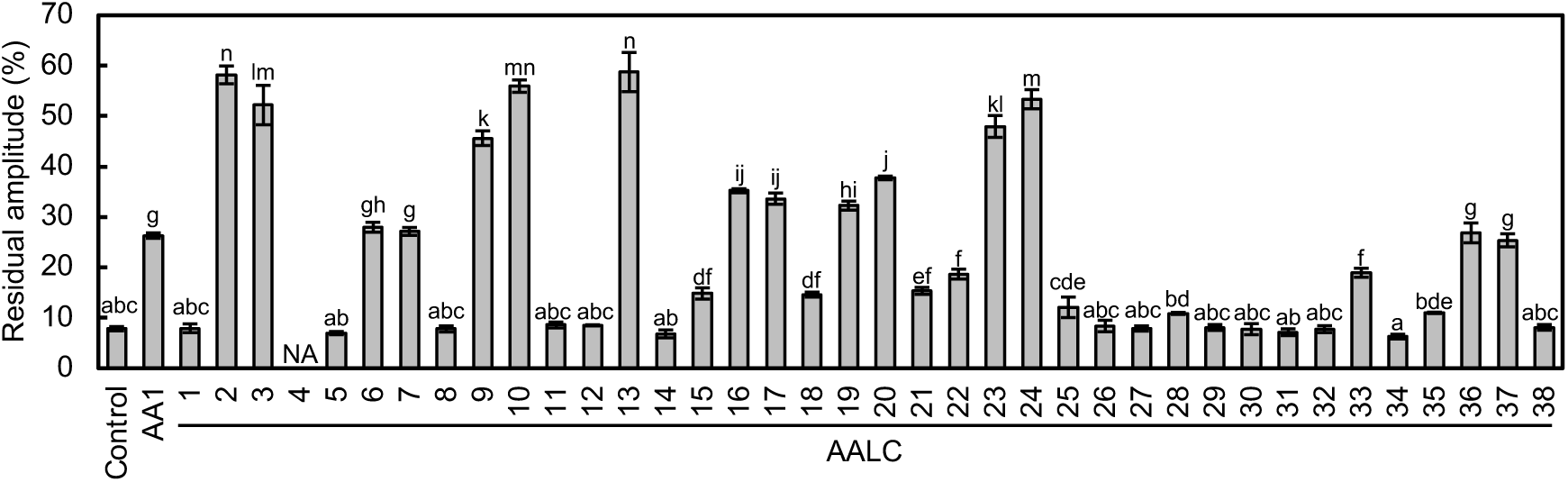
Effects of AALCs on the residual component of Q_A_^−^ reoxidation kinetics in the presence of DCMU. The % amplitudes of the residual components obtained from the analysis of Q_A_^−^ reoxidation measurements in the presence of DCMU are shown. Data are mean ± SD (*n* = 3). Different lower-case letters above bars indicate statistically significant differences (*P* < 0.05, Tukey’s HSD test).

Previous SAR studies of AALCs on complex III showed a tendency that the inhibitory activity rises as the acidity of the phenolic OH group, especially important for the inhibitory activity, increases (Neft and Farley, 1971; Rieske, 1976; Tokutake et al., 1994). This appears to apply to the inhibitory activities of AALCs on PSII, as well. None of the compounds without nitro or formylamino substituents on the salicylic acid moiety showed effects on PSII in this assay. This suggests that the nitro- or formylamino-substituted salicylic acid moiety is crucial for the AA effect on PSII. A common feature among AALCs with effects stronger than AA1 is the presence of a nitro group, which has a strong electron-withdrawing property, at position 3 or 5 in the salicylic acid moiety. Both the 3-nitro-substituted AALC2 and 5-nitro-substituted AALC3 showed very strong effects on PSII. Compared to AALC2, AALC9 with an additional 5-methyl substituent, which has an electron-donating property, showed decreased inhibitory activity. AALC6 with 4-methyl-3-nitro substitution further decreased the inhibitory activity. This could be because the presence of a vicinal methyl decreases the electron-withdrawing property of the nitro group, since the steric congestion arising from vicinal methyl leads to a twisted plane of the nitro group against the benzene-ring plane (Tokutake et al., 1994). The 4-methyl substituent might also sterically hinder AALC-binding to PSII. The decreased inhibitory activity of AALC7 compared to AALC3 can be explained in the same way. The reason behind AALC10 having the same activity as AALC3 is likely to be more complex, but a similar tendency was observed previously when inhibition of complex III was studied (Tokutake et al., 1994). In contrast to AALC9, AALC13, which has a 5-chloro substituent (a weak electron-withdrawing group) in addition to a 3-nitro substituent, showed strong inhibitory activity similar to that of AALC2. The presence of only 5-chloro- (AALC12) or 5-bromo-substitution (AALC11), which have weak electron withdrawing properties, was insufficient to exhibit inhibitory effects. However, 5-chloro-substitution crucially supported the inhibitory activity of the 3-formylamino-subtituted AALC33; other 3-formylamino-substituted AALCs with dialkyl amino acid esters instead of the substituted dilactone ring did not exhibit inhibitory activity. It should be noted that, although the formylamino group itself has a poor electron withdrawing property, formylamino substitution in *ortho* position to the phenolic OH has been suggested to decrease the p*K*_a_ of the phenolic OH by *ortho* effects (proximity effects) (Tokutake et al., 1994).

The amide linkage between the salicylic acid moiety and the hydrophobic structure also seems to play an important role in the inhibitory activity of AALCs. While AALC2 with a 3-nitrosalicylic acid (2-hydroxy-3-nitrobenzoic acid) moiety showed a strong effect, AALC26 with a 2-hydroxy-3-nitrophenylacetic acid moiety, in which the amide linkage and the phenolic benzene ring are separated by a methylene group, had no effects. In AALC2, but not in AALC26, the phenolic OH group can form an intramolecular hydrogen bond (H-bond) with either the carbonyl O or the NH of the amide linkage. The former intramolecular H-bond has been shown to be formed in free AA/AALCs (Miyoshi et al., 1995; Kim et al., 1999), whereas the latter (with NH as the H-bond donor) has been suggested to be formed in AA bound to complex III (Huang et al., 2005). The formation of an intramolecular H-bond between the phenolic OH group and the amide linkage, required for AA’s inhibitory activity against complex III (Miyoshi et al., 1995), seems to be important for its activity against PSII, as well. This is consistent with the observation that the *N*-methylation of the amide linkage leads to a substantial loss of inhibitory activity, as suggested from comparison between AALC37 and AALC38 (and partially, between AALC24 and AALC25). The NH of the amide linkage could also be directly involved in the interaction between AA/AALCs and PSII. It should be noted that AALC15, widely known as dinoseb, showed a small but significant effect, even though it lacked the amide group. If dinoseb binds to the same site as AA in PSII, this would indicate that the amide group may not be absolutely essential, although generally required.

While the nitro- or formylamino-substituted salicylamide (2-hydroxybenzamide) structure is crucial for the inhibitory activity, the natural dilactone-ring structure of AA1 is replaceable by various flexible hydrophobic structures. However, the presence of a hydrophobic structure is necessary, as indicated by the abolished activity of AALC14. When the salicylic acid moiety is nitro-substituted, the restrictions on the hydrophobic structures seem to be weak. Comparisons among nitrosalicylamide derivatives that have various dialkyl glutamate or aspartate esters (AALC2, 3, and 16–21) suggest that there is an optimal length of alkyl chains (or optimal hydrophobicity); the inhibitory activity measured as the residual amplitude was found to be C5 (14.6–32.2%) < C9 (33.6–37.7%) < C8 (52.2–58.2%). A similar tendency was observed in previous studies on the complex III–inhibiting activity of various formylaminosalicyalmide derivatives (Tokutake et al., 1993). In this assay, we did not observe differences between the activity of AALC16 derived from L-form aspartate and AALC17 derived from D-form aspartate. The reason for the notable difference of activity between AALC18 (3-nitro) and AALC19 (5-nitro) is unclear. Nitrosalicylamide derivatives with substituted diphenyl ether groups (AALC23 and 24) showed strong inhibitory activities only slightly lower than that of AALC2. When the salicylic acid moiety is formylamino-substituted, like in natural AA, the dependence of the inhibitory activity on the hydrophobic structure increased. Unless there was a chloro group in addition to the formylamino group (AALC33), neither various dialkyl glutamate esters (AALC27–29), an *n*-pentyl group (AALC30), nor a phenyl group (AALC34) were able to replace the natural dilactone ring moiety of AA1. Moreover, while AALC35 with a diphenyl ether group did not show inhibitory activity on PSII, *tert*-butyl substitution (AALC36) or di(*sec*-butyl) substitution (AALC37) of the diphenyl ether group led to a notable inhibitory activity similar to that of AA1. This is similar to the difference between AA3 and AA1, considering that the inhibitory effects on PSII observed for AA1 are abolished in AA3, in which the *n*-hexyl substituent in the dilactone ring moiety is replaced by a shorter *n*-butyl group.

### 3.2. *F*_v_/*F*_m_ of PSII treated with AA-like compounds

Natural AA does not affect the initial *F*_v_/*F*_m_ of PSII in the dark (Imaizumi et al., 2025). However, given that some of the AALCs, such as AALC4, exhibited particularly pronounced effects on PSII, we examined whether the AALCs affected the initial *F*_v_/*F*_m_ in the absence of DCMU (**Fig. 4**). The *F*_v_/*F*_m_ in thylakoid membranes without inhibitors was 0.809, and treatment with AA1 did not have significant effects on the *F*_v_/*F*_m_, consistent with our previous report. While most AALCs did not affect the initial *F*_v_/*F*_m_, addition of AALC4 led to a remarkable drop to *F*_v_/*F*_m_ = 0.411. In addition, AALC2, 24, and 25 induced a decrease in the initial *F*_v_/*F*_m_ to below 0.7 (AALC2, *F*_v_/*F*_m_ = 0.675; AALC24, *F*_v_/*F*_m_ = 0.665; AALC25, *F*_v_/*F*_m_ = 0.588), and a slight decrease was observed with AALC3, 9, 13, 16, 17, and 23 (*F*_v_/*F*_m_ = 0.70–0.76). All of these AALCs possess a large hydrophobic moiety and a 3-nitrosalicylic acid–derived moiety, except for AALC3, which contains a 5-nitrosalicylic acid moiety. The decrease of the initial *F*_v_/*F*_m_ was due to a notable decline in the *F*_m_ level; none of the AALCs mentioned above induced a rise in the *F*_o_ level.

**Fig. 4.**
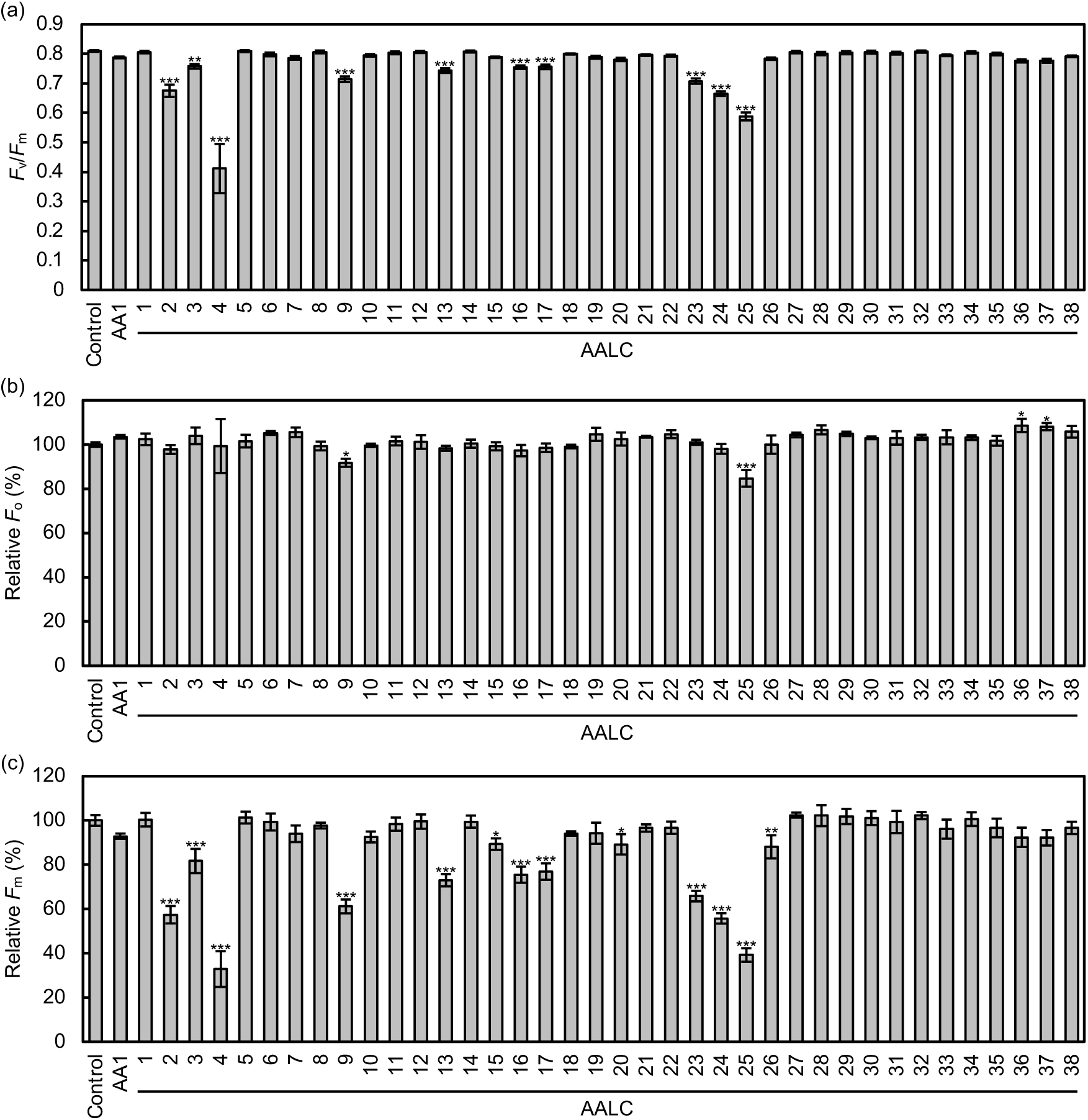
Effects of AALCs on the initial *F*_v_/*F*_m_ values. The (**a**) *F*_v_/*F*_m_ values, (**b**) relative *F*_o_ levels (% of control), and (**c**) relative *F*_m_ levels (% of control) of thylakoid membranes treated with ethanol (control), 10 µM AA1, or 10 µM AALCs in the dark are shown. Data are mean ± SD (*n* = 3). Asterisks above bars indicate statistically significant differences from the control (\*\*\**P* < 0.001, \*\**P* < 0.01, \**P* < 0.05, Dunnett’s test).

### 3.3. Effects of AA-like compounds on the Q_A_^−^ reoxidation kinetics in the absence of DCMU

AA alone (without DCMU) was previously found to delay the reoxidation of Q_A_^−^ (Takagi et al., 2019; Imaizumi et al., 2025). In the absence of DCMU, the decay curve can be fitted to three exponential components and a time-independent residual component, and the fast decay component represents electron transfer from Q_A_^−^ to Q_B_, the intermediate decay component reflects electron transfer from Q_A_^−^ to Q_B_ after the binding of a PQ molecule to an empty Q_B_ site, the slow decay component represents charge recombination of Q_A_^−^ with the donor side of PSII, and the residual component may be attributed to the equilibrium between Q_A_^−^Q_B_ and Q_A_Q_B_^−^ (Reifarth et al., 1997). In the presence of AA, the proportion of the fast phase decreased greatly, and the time constants for the fast and intermediate phases increased significantly, suggesting that AA suppresses electron transfer from Q_A_^−^ to Q_B_ and the binding of PQ to an empty Q_B_ site (Takagi et al., 2019; Imaizumi et al., 2025).

To further study the effects of AALCs on PSII, we conducted Q_A_^−^ reoxidation measurements of thylakoid membranes in the absence of DCMU (**Fig. 5** and **Table 1**). AA1 notably diminished the amplitude of the fast phase and tripled the time constants for the fast and intermediate phases, suggesting the suppression of Q_A_^−^-to-Q_B_ electron transfer and PQ binding to an empty Q_B_ site, consistent with our previous results. All AALCs that showed effects in the presence of DCMU, suppressed Q_A_^−^ reoxidation to some extent in the absence of DCMU, as well (except for AALC33, which only showed a slight effect in the absence of DCMU). Although to different extents, these compounds decreased the fast phase amplitude and increased the time constants for the fast and intermediate phases, as was the case with AA1. AALC4 showed strong effects on PSII, but again we were unable to obtain a reliable decay curve due to a low signal-to-noise ratio (**Fig. S2b**). Although similar tendencies were observed, there were some differences between the SAR in the absence and presence of DCMU, and the SAR in the absence of DCMU was less clear.

**Fig. 5.**
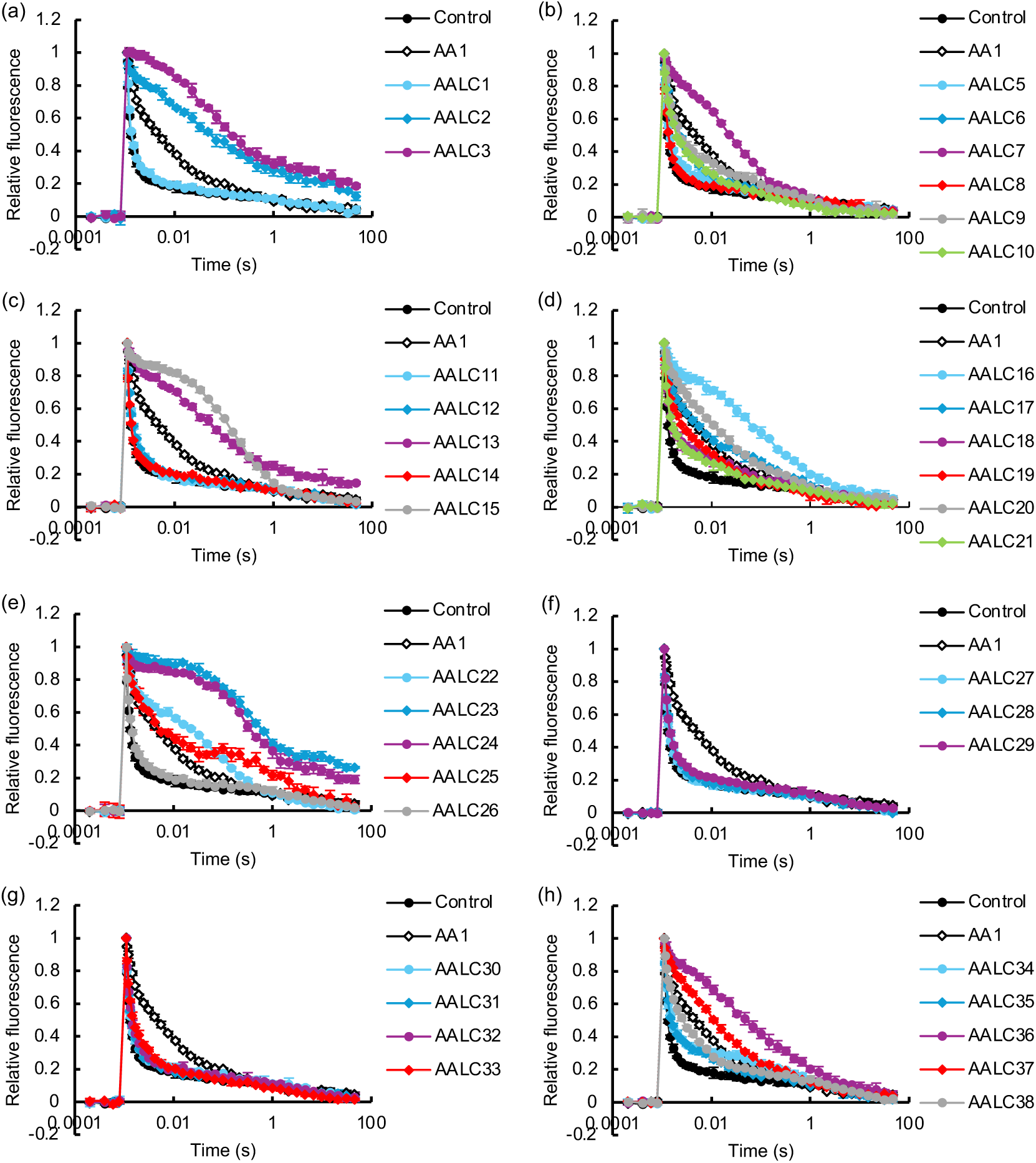
Effects of AALCs on the Q_A_^−^ reoxidation kinetics of PSII (in the absence of DCMU). Ethanol (“control”; black filled circles), 10 µM AA1 (black open diamonds), and 10 µM AALCs (colored filled symbols) were added to spinach thylakoid membranes. Although all compounds (and the control) were measured in each independent experiment, for clarity, the results for AALCs are presented in eight separate panels (**a**–**h**), with the same control and AA1-treated data shown in all panels. Data are mean ± SD (*n* = 3). In some instances, error bars are smaller than the symbols.

**Table 1.** Kinetic parameters of Q_A_^−^ reoxidation in thylakoid membranes treated with ethanol (control), 10 µM AA1, or 10 µM AALCs.

| Compound | Fast phase |  | Intermediate phase |  | Slow phase |  | Residual |
| --- | --- | --- | --- | --- | --- | --- | --- |
| | A1 (%) | τ1 (ms) | A2 (%) | τ2 (ms) | A3 (%) | τ3 (s) | |
| Control | 78.1 | 0.15 | 12.3 | 2.7 | 6.0 | 1.6 | 3.6 |
| AA1 | 41.3 | 0.48 | 37.5 | 10.2 | 15.4 | 0.5 | 5.8 |
| AALC1 | 75.7 | 0.17 | 13.7 | 3.5 | 6.8 | 1.4 | 3.8 |
| AALC2 | 31.3 | 5.09 | 36.9 | 144.6 | 16.7 | 11.9 | 15.1 |
| AALC3 | 25.9 | 15.05 | 39.8 | 16.0 | 14.8 | 10.1 | 19.5 |
| AALC5 | 71.6 | 0.18 | 16.4 | 4.6 | 7.4 | 0.8 | 4.6 |
| AALC6 | 47.8 | 0.35 | 34.6 | 4.2 | 13.8 | 0.8 | 3.8 |
| AALC7 | 21.1 | 1.11 | 54.5 | 36.0 | 19.1 | 1.1 | 5.3 |
| AALC8 | 69.9 | 0.11 | 19.6 | 1.1 | 6.6 | 0.4 | 3.9 |
| AALC9 | 57.4 | 0.48 | 21.6 | 12.6 | 16.8 | 0.8 | 4.2 |
| AALC10 | 54.8 | 0.28 | 28.3 | 5.2 | 13.8 | 0.4 | 3.1 |
| AALC11 | 66.4 | 0.17 | 23.1 | 2.1 | 7.6 | 2.5 | 2.9 |
| AALC12 | 67.7 | 0.16 | 20.7 | 2.2 | 8.1 | 1.3 | 3.5 |
| AALC13 | 21.6 | 0.98 | 41.2 | 48.2 | 20.5 | 1.1 | 16.7 |
| AALC14 | 78.3 | 0.15 | 11.1 | 4.2 | 7.2 | 1.4 | 3.3 |
| AALC15 | 14.1 | 0.36 | 42.3 | 87.2 | 38.1 | 0.7 | 5.5 |
| AALC16 | 21.6 | 0.55 | 37.7 | 47.8 | 32.2 | 1.1 | 8.6 |
| AALC17 | 38.7 | 0.33 | 31.7 | 7.2 | 25.4 | 0.7 | 4.2 |
| AALC18 | 68.0 | 0.26 | 17.5 | 17.8 | 11.9 | 2.0 | 2.7 |
| AALC19 | 53.9 | 0.36 | 31.0 | 12.4 | 12.7 | 0.9 | 2.4 |
| AALC20 | 31.9 | 0.80 | 40.7 | 19.4 | 21.2 | 0.7 | 6.3 |
| AALC21 | 67.0 | 0.27 | 19.4 | 11.6 | 10.8 | 1.5 | 2.9 |
| AALC22 | 37.5 | 0.44 | 38.6 | 53.7 | 20.6 | 1.0 | 3.3 |
| AALC23 | 8.0 | 1.13 | 53.6 | 372.1 | 12.3 | 15.7 | 26.1 |
| AALC24 | 15.1 | 0.29 | 51.5 | 307.3 | 14.4 | 7.9 | 19.0 |
| AALC25 | 35.5 | 0.32 | 31.4 | 5.8 | 27.6 | 2.1 | 5.5 |
| AALC26 | 71.0 | 0.18 | 17.2 | 3.0 | 8.9 | 2.8 | 2.9 |
| AALC27 | 70.9 | 0.15 | 19.2 | 1.8 | 7.5 | 2.2 | 2.4 |
| AALC28 | 67.5 | 0.19 | 21.5 | 2.3 | 8.9 | 2.4 | 2.1 |
| AALC29 | 70.0 | 0.19 | 17.2 | 3.1 | 9.5 | 1.8 | 3.3 |
| AALC30 | 75.6 | 0.16 | 13.3 | 3.8 | 8.0 | 2.5 | 3.1 |
| AALC31 | 72.9 | 0.18 | 16.5 | 3.2 | 7.4 | 1.9 | 3.1 |
| AALC32 | 67.6 | 0.14 | 20.9 | 2.0 | 8.6 | 1.8 | 3.0 |
| AALC33 | 64.6 | 0.19 | 23.8 | 3.3 | 9.3 | 1.1 | 2.3 |
| AALC34 | 66.3 | 0.16 | 14.2 | 2.7 | 15.6 | 0.9 | 4.0 |
| AALC35 | 68.6 | 0.23 | 18.5 | 11.6 | 10.6 | 2.8 | 2.3 |
| AALC36 | 18.8 | 0.52 | 41.7 | 32.4 | 31.1 | 1.1 | 8.4 |
| AALC37 | 27.6 | 0.76 | 45.7 | 17.6 | 21.1 | 0.8 | 5.6 |
| AALC38 | 47.2 | 0.32 | 35.5 | 6.4 | 14.0 | 2.9 | 3.3 |

As was the case in the presence of DCMU, none of the compounds without nitro or formylamino substituents on the salicylic acid moiety showed clear effects on PSII. In contrast, all 3- and 5-nitrosalycilamide derivatives (excluding AALC14, which lacks a hydrophobic structure replacing the natural dilactone ring) and some 3-formylaminosalycilamide derivatives showed inhibitory effects, reconfirming the critical role of the salicylamide structure. Considering that the inhibitory effects of AALC3, 7, and 19 tended to be stronger than that of AALC2, 6, and 18, respectively, 5-nitro substitution may be slightly more favorable than 3-nitro substitution in terms of inhibitory activity in the absence of DCMU.

AALC2, 3, 13, 23, and 24 had strong effects both in the presence (residual component above 47%) and absence (fast phase amplitudes between 8% and 32%) of DCMU. These AALCs showed a clear rise of the residual component even in the absence of DCMU (amplitudes above 15%), and may possess both AA-like properties and weak DCMU-like properties. On the other hand, while AALC9 and 10 also showed much stronger effects than AA1 in the presence of DCMU, their effects in the absence of DCMU were slightly weaker than AA1, but the underlying reason is unclear. In contrast, AALC15 (dinoseb) showed a very strong effect (fast phase amplitude of 14.1%) in the absence of DCMU, although its effect in the presence of DCMU was weaker than AA1 (residual amplitude of 14.8%). It could be possible that dinoseb can bind to or near the Q_B_ site, in addition to the AA-binding site.

The effect of the amide linkage between the salicylic acid moiety and the hydrophobic structure on the inhibitory activity of AALCs was also observed in the absence of DCMU, but its impact was weaker than under the presence of DCMU. In contrast to AALC2 showing a strong effect, AALC26, whose amide linkage and the phenolic benzene ring are separated by a methylene group, showed little effect on Q_A_^−^ reoxidation. This suggests the importance of the intramolecular H-bond between the phenolic OH group and the amide linkage, as discussed earlier. However, comparison between AALC37 and AALC38 (and partially, between AALC24 and AALC25) shows that *N*-methylation of the amide linkage only partially decreases the inhibitory effect of the AALCs. Taking the results under the presence of DCMU into consideration, the current results would indicate that, in terms of inhibitory activity in the absence of DCMU, the required intramolecular H-bond between the phenolic OH group and the amide linkage does not need to involve the NH of the amide linkage, and that the NH of the amide linkage is not involved in critical interactions between AA/AALCs and PSII.

Similar to the case in the presence of DCMU, the dilactone ring moiety of AA1 was replaceable by various hydrophobic structures. Comparison among AALC3, 19, and 20 suggested that, also in the absence of DCMU, there is a similar optimal length for the alkyl chains (or optimal hydrophobicity). The inhibitory activity of formylamino-substituted AALCs was more dependent on the hydrophobic structure, as was also the case under the presence of DCMU. AALCs with dialkyl glutamate esters or an *n*-pentyl group replacing the natural dilactone ring moiety of AA1 (AALC27–33) showed no or only slight effects. While AALC35 with a diphenyl ether group only showed weak effects on PSII (fast phase amplitude of 68.6%), AALC36 with a *tert*-butyl substitution of the diphenyl ether group showed remarkable inhibitory activity (fast phase amplitude of 18.8%), again resembling the difference between AA3 and AA1.

## 4. Discussion

In this study, we compared the effects of various AALCs on PSII. The results show that AALCs with certain structural features exert AA-like effects on PSII to varying extents. Substituents on the salicylic acid moiety that increase the acidity of the phenolic OH and the presence of a free NH–containing amide linkage adjacent to the phenolic ring were especially important for inhibitory activity against PSII, similar to when inhibiting complex III (Rieske, 1976; Tokutake et al., 1994; Miyoshi et al., 1995). The hydrophobic structure corresponding to the dilactone ring moiety in AA also affected the inhibitory activity, but it did not need to be a dilactone ring, and was replaceable by a variety of structures.

The acidity of the phenolic OH was important for the inhibitory activity. Also, it was essential that the amide linkage was present adjacent to the phenolic ring. These observations resemble the results of SAR studies on the binding of AALCs to complex III (Rieske, 1976; Tokutake et al., 1994; Miyoshi et al., 1995), and suggest that (1) the phenolic OH forms a H-bond with PSII, and that (2) a critical intramolecular H-bond between the phenolic OH and the amide linkage is formed, as was the case in the binding of AA to complex III (Huang et al., 2005). While the overall SAR trends of AALCs in PSII were similar to those observed in complex III, one major difference emerged: in complex III, AALCs bearing a 3-formylamino substituent on the salicylic acid moiety exhibited higher inhibitory activity than those with a nitro substituent (Tokutake et al., 1994), whereas in PSII, those with 3- or 5-nitro substituents were more active than those with a 3-formylamino substituent. Crystal structures and cryo-electron microscopy structures of complex III with AA bound revealed that both the phenolic OH and the formylamino NH of the 3-formylaminosalicylic acid moiety of AA form H-bonds with an acidic residue at the Q_i_ site (Asp228 of Cyt *b* in bovine sequence) (Huang et al., 2005; Esser et al., 2008; Wieferig and Kühlbrandt, 2023). This suggests that the preference for the 3-formylamino group over a nitro group in AALCs when binding to complex III, was likely due to the stabilizing effect of the additional H-bond formed between the formylamino group and the Q_i_ site. Conversely, the weaker effect of the 3-formylamino group on the binding of AA to PSII suggests that it may or may not form a H-bond with PSII, and even if formed, its contribution to binding is not substantial.

Various hydrophobic structures were able to replace the dilactone ring moiety in nitrosalicylamide-derived AALCs. There did, however, seem to be an optimal hydrophobicity, and inhibitory activity was not observed without the hydrophobic moiety. Meanwhile, stronger structural constraints were imposed on the hydrophobic moiety in formylaminosalicylamide-derived AALCs. This suggests that the hydrophobic moiety is required to increase the hydrophobicity of AALCs, most likely to enter hydrophobic cavities in PSII, and that it may also facilitate effective interaction of the formylamino-substituted salicylic acid moiety with the binding pocket. There may additionally be a specific important hydrophobic interaction between the inhibitor and a PSII component, causing the critical difference between the effects of AA1 and AA3 on PSII (Imaizumi et al., 2025; Imaizumi and Ifuku, 2025b), but further structural studies are required to address this point.

Based on our current results, our previous results on different AA components (Imaizumi et al., 2025), and comparison with previous SAR studies of AA-like compounds on complex III (Tokutake et al., 1993; Tokutake et al., 1994; Miyoshi et al., 1995), we propose two tentative models for the binding mode of AA (AA1) to PSII (**Fig. S3**). The phenolic OH of AA1 is likely to be involved in the H-bond formation with PSII, as well as in the formation of a six-membered intramolecular H-bond with either (1) the amide NH (**Fig. S3a**) or (2) the amide carbonyl (**Fig. S3b**). If it is the latter case, the amide NH may be directly involved in an interaction, presumably an H-bond, with PSII. If the phenolic OH of AA1 is deprotonated, AA1 could bind to PSII in a similar manner as the former model (**Fig. S3a**), but with its deprotonated phenolic OH serving as an H-bond acceptor, and an amino acid side chain serving as an H-bond donor. In contrast to its binding mode to complex III, it is unclear whether the formylamino group is directly involved in interaction with PSII. The dilactone ring moiety increases the hydrophobicity of the molecule and may support the tight fitting of AA1 into its binding cavity by regulating the whole molecular configuration together with the amide linkage. Considering the presence and absence of inhibitory effects of AA1 and AA3, respectively (Imaizumi et al., 2025; Imaizumi and Ifuku, 2025b), the *n*-hexyl chain of the natural dilactone ring moiety might also be involved in hydrophobic interactions, which may be crucial when the salicylic acid moiety contains the natural 3-formylamino group.

Although there was a similar trend between the structure–activity relationship in the presence and absence of DCMU, the results did not completely match with each other (**Fig. 2**, **Fig. 3**, **Fig. 5**, and **Table 1**). This, however, was as expected, since previous Q_A_^−^ reoxidation measurements showed that AA1 has a stronger effect than AA2 in the presence of DCMU, whereas AA2 shows a slightly stronger effect than AA1 in the absence of DCMU (Imaizumi et al., 2025). These observations suggest that the detailed mechanisms underlying the AA-effects in the presence and absence of DCMU are not exactly the same. We suspect that even at the same binding site, differences in the mode of interaction with PSII can modulate the effects of AALCs on PSII. While the AA-effects were suggested to reach near-saturation at 10 µM AA (Imaizumi et al., 2025), the increase of the residual amplitude in the presence of DCMU by some AALCs (i.e. AALC2, 3, 9, 10, 13, 23, and 24) at 10 µM were remarkably higher than that by 10 µM AA1 (**Fig. 2** and **Fig. 3**). This also supports that the variations in the AA-effects of AALCs most likely cannot be explained solely by their binding affinities to PSII; instead, it would also reflect differences in their detailed interaction with PSII. As exceptions, some of the nitrosalicylamide-derived AALCs that, unlike AA, induced a significant light-independent drop in *F*_v_/*F*_m_ (**Fig. 4**) may have additionally perturbed a distinct region in PSII.

AALC15 (dinoseb), a well-known phenolic PSII inhibitor (Fuerst and Norman, 1991), had effects similar to those of DCMU, while it also exhibited some weak AA-like effects. Previous studies have shown that dinoseb converts high-potential Cyt *b*_559_ to its intermediate- or low-potential form (Rutherford et al., 1984; Kaminskaya et al., 2007) and that it could act as an ADRY (acceleration of the deactivation reactions of the water-splitting enzyme system Y) reagent (Rutherford et al., 1984); such properties have also been suggested for AA (Cramer et al., 1971; Yerkes and Crofts, 1992; Takagi et al., 2019). Dinoseb has been suggested to bind near the acceptor side of PSII, but possibly to a quinone-binding site near Cyt *b*_559_ (Q_C_ or Q_D_ site) rather than or in addition to the Q_B_ site (DCMU-binding site) (Oettmeier and Masson, 1980; Kaminskaya et al., 2007; Kaminskaya and Shuvalov, 2013; Yadav et al., 2014). It should be noted that these “Q_C_” and “Q_D_” sites do not necessarily correspond to the Q_C_ and Q_D_ sites identified in structural studies of PSII (Guskov et al., 2009; Kamada et al., 2023). It could be possible that, like dinoseb, AA1 also binds to a potential quinone-binding site near the PSII acceptor side and Cyt *b*_559_, as speculated previously (Imaizumi et al., 2025).

The effects of two AA-like compounds on PGR5-dependent CEF-PSI have been investigated in a previous study through Fd-dependent PQ reduction assays (Taira et al., 2013). The compound they refer to as “AAL1” appears to be identical to our AALC20. It should be noted, however, that the stereochemistry of the chiral carbon adjacent to the amide bond in their figure seems inverted. AALC20, or “AAL1,” inhibited PGR5-dependent CEF-PSI more efficiently than AA3, while seemingly targeting the same inhibitory site. The other compound, which they refer to as “AAL2,” is similar to our AALC2 and 16 (or 17), and was also found to be more potent than AA3 at inhibiting PGR5-dependent CEF-PSI. In our current study, we have shown that these compounds also have stronger inhibitory effects on PSII than AA1 does (**Fig. 2**, **Fig. 3**, **Fig. 5**, and **Table 1**). These results suggest that, at present, it may be challenging to inhibit PGR5-dependent CEF-PSI without impacting PSII, except through the use of AA3. Even though the target of AA in inhibiting PGR5-dependent CEF-PSI is expected to be distinct from PSII (Sugimoto et al., 2013; Imaizumi et al., 2025; Imaizumi and Ifuku, 2025b), some structural features of AA required for the inhibition of the PGR5-dependent pathway and for the inhibitory effects on PSII may be similar.

Although the molecular mechanism remains to be elucidated, some of the AALCs exhibited especially pronounced effects on PSII. As we have demonstrated earlier, the rise of the residual component in Q_A_^−^ reoxidation measurements is indicative of a drop in *F*_v_/*F*_m_ due to increased *F*_o_ by a single-turnover flash (Imaizumi et al., 2025). Several AALCs induced a remarkable rise of the residual component in the presence of DCMU (**Fig. 3**), and some of them also increased this component even in the absence of DCMU (**Fig. 5** and **Table 1**). There were also AALCs that significantly decreased *F*_v_/*F*_m_ even before exposure to any light (**Fig. 4**). Such effect is not universally observed among phenolic PSII inhibitors, as evident from the results with dinoseb (AALC15) in the current study and bromoxynil in our previous report (Imaizumi et al., 2025). While it is unclear whether the mechanism is the same, 3-nitro-2,4,6-trihydroxybenzamide derivatives, similar to our nitrosalicyalmide-derived AALCs, but with two additional hydroxy groups, have also been reported to be effective inhibitors of photosynthetic electron transport (Honda et al., 1990a; Honda et al., 1990b). Therefore, potent AA-like PSII inhibitors found in our current study could potentially serve as lead compounds for the development of new herbicides.

## CRediT authorship contribution statement

**Ko Imaizumi:** Writing – original draft, Writing – review & editing, Visualization, Validation, Methodology, Investigation, Funding acquisition, Formal analysis, Data curation. **Masatoshi Murai:** Writing – review & editing, Resources. **Hideto Miyoshi:** Writing – review & editing, Resources. **Kentaro Ifuku:** Writing – review & editing, Supervision, Resources, Project administration, Funding acquisition, Conceptualization.

## Data Availability

The data underlying this article are available in the article and its online supplementary data.

