## supplementary data for "Structure–activity relationship of antimycin A-like compounds as photosystem II inhibitors"

**Contents:**

**Fig. S1** Chemical structures of natural AA components.
**Fig. S2** Effects of AALC4 on the $Q_A^-$ reoxidation kinetics of PSII.
**Fig. S3** Two tentative models for AA1 binding in its binding cavity in PSII.

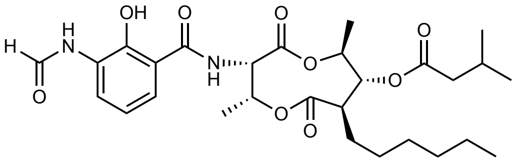

AA1

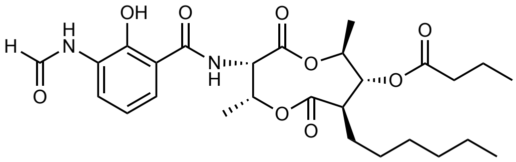

AA2

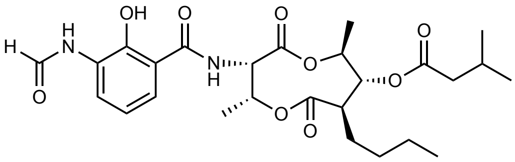

AA3

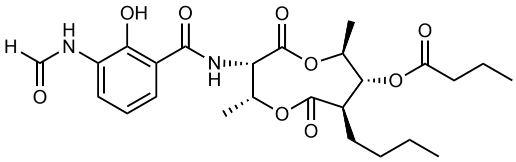

AA4

**Fig. S1** Chemical structures of natural AA components. The structures of AA1, AA2, AA3, and AA4 are shown.

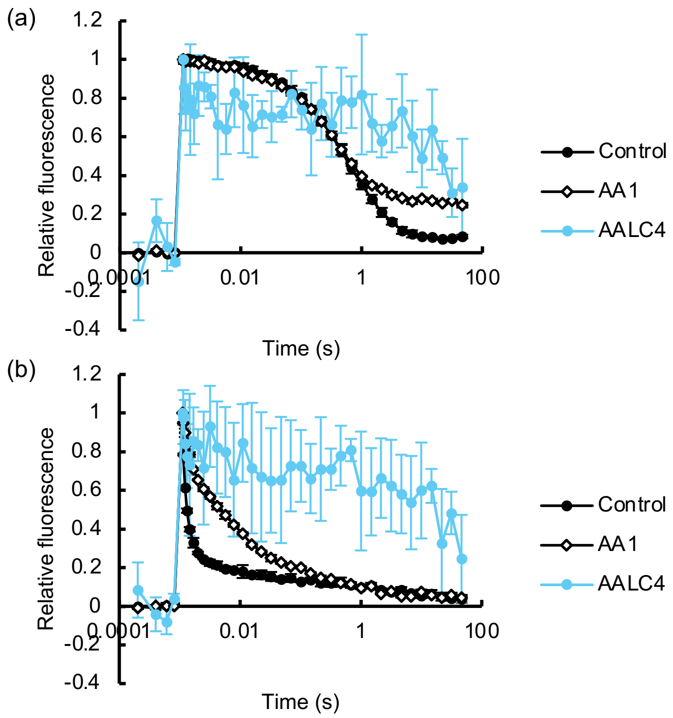


**Fig. S2** Effects of AALC4 on the $Q_A^-$ reoxidation kinetics of PSII. Ethanol ("control"; black filled circles), 10 µM AA1 (black open diamonds), and 10 µM AALC4 (blue filled circles) were added (**a**) together with or (**b**) without 10 µM DCMU to spinach thylakoid membranes. Although shown here for clarity, the measurements for AALC4 were conducted together with the control and all other compounds in Fig. 2 (for panel **a**) and Fig. 5 (for panel **b**); thus, the control and AA1-treated data are the same as those in Fig. 2 (for panel **a**) and Fig. 5 (for panel **b**). Data are mean ± SD (*n* = 3). In some instances, error bars are smaller than the symbols.

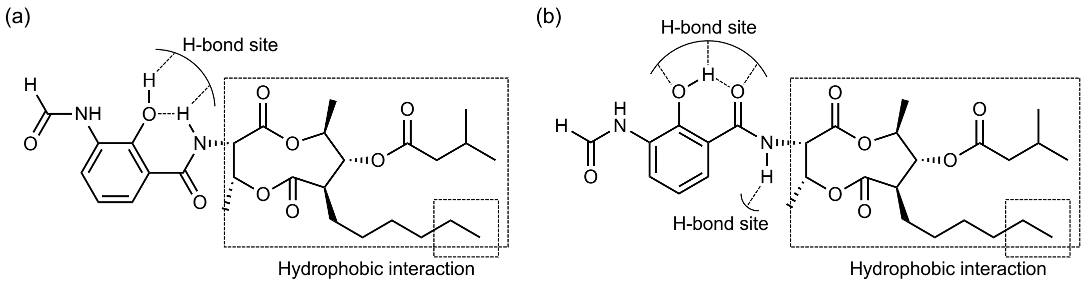


**Fig. S3** Two tentative models for AA1 binding in its binding cavity in PSII. The phenolic OH is expected to form an intramolecular H-bond with (**a**) the amide NH or (**b**) the amide carbonyl.
